# A new parallel high-pressure packing system enables rapid multiplexed production of capillary columns

**DOI:** 10.1101/2021.02.26.433033

**Authors:** Johannes B. Müller-Reif, Fynn M. Hansen, Lisa Schweizer, Peter V. Treit, Philipp E. Geyer, Matthias Mann

**Affiliations:** Department of Proteomics and Signal Transduction, Max Planck Institute of Biochemistry, Martinsried, Germany; NNF Center for Protein Research, Faculty of Health Sciences, University of Copenhagen, Copenhagen, Denmark; OmicEra Diagnostics GmbH, Planegg, Germany

## Abstract

Reversed-phase high performance liquid chromatography (HPLC) is the most commonly applied peptide separation technique in mass spectrometry (MS)-based proteomics. Particle-packed capillary columns are predominantly used in nano-flow HPLC systems. Despite being the broadly applied standard for many years capillary columns are still expensive and suffer from short lifetimes, particularly in combination with ultra-high-pressure chromatography systems. For this reason, and to achieve maximum performance, many laboratories produce their own in-house packed columns. This typically requires a considerable amount of time and trained personnel. Here, we present a new packing system for capillary columns enabling rapid, multiplexed column production with pressures reaching up to 3000 bar. Requiring only a conventional gas pressure supply and methanol as driving fluid, our system replaces the traditional setup of helium pressured packing bombs. By using 10x multiplexing, we have reduced the production time to just under 2 minutes for several 50 cm columns with 1.9 µm particle size, speeding up the process of column production 40 to 800 times. We compare capillary columns with various inner diameters (ID) and length packed under different pressure conditions with our newly designed, broadly accessible high-pressure packing station.

**One sentence summary:** A newly constructed parallel high-pressure packing system enables the rapid multiplexed production of capillary columns.

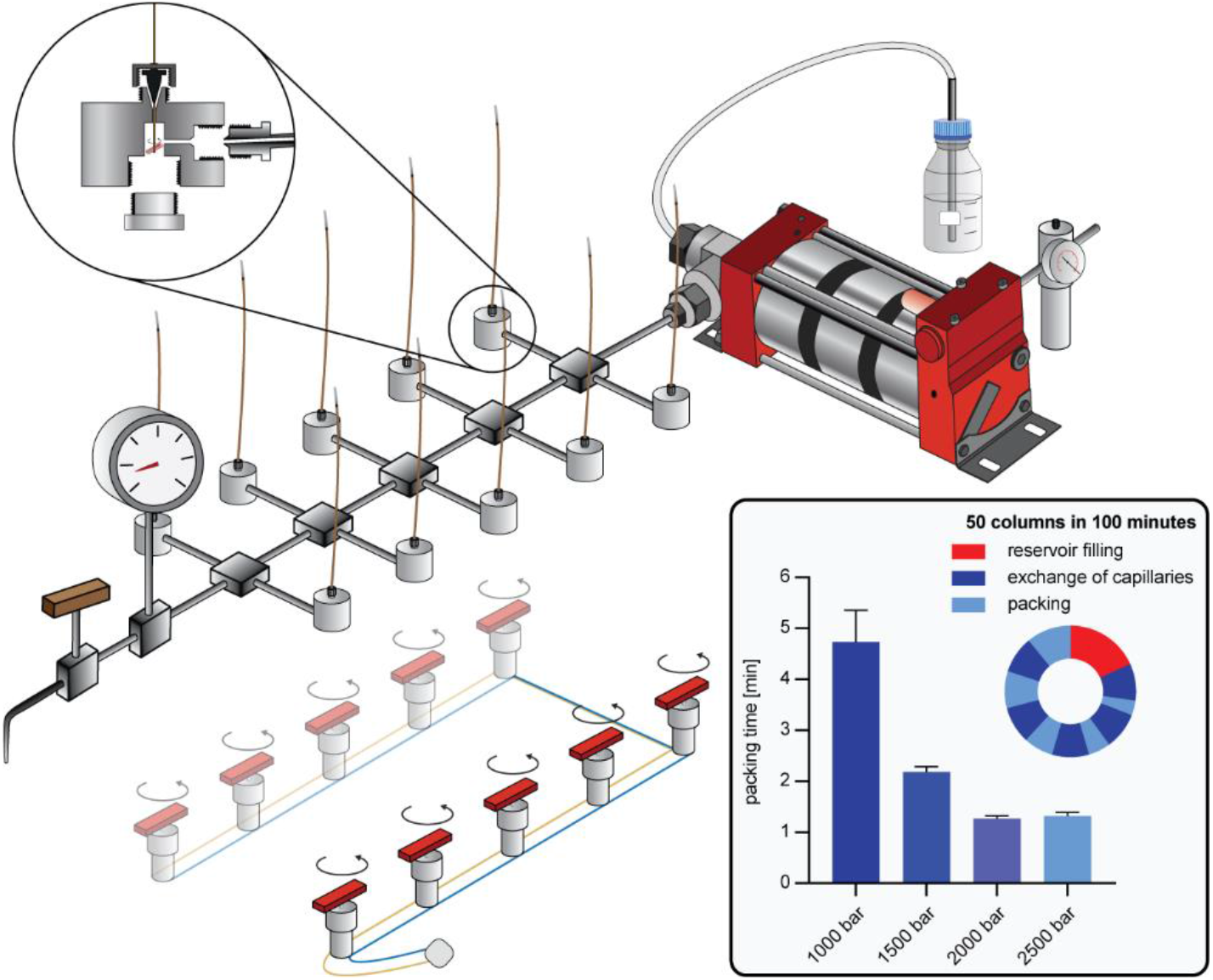

## Introduction

State-of-the-art mass spectrometry (MS)-based proteomic pipelines typically consist of a sample preparation workflow to digest proteins and harvest pure peptides, a liquid chromatography (LC) system for peptide separation, a mass spectrometer and a sophisticated bioinformatics pipeline for raw data interpretation and subsequent statistical analysis (1,2). The LC system plays a central role by separating the complex mixture of tens of thousands of peptides in a time-resolved manner according to their biochemical properties, making them ultimately manageable for the MS system over the course of a gradient (3,4). The most widely applied technique for high-performance applications is reversed-phase separation, originally introduced in the 1970s (5). In essence, chromatographic systems are made of programmable pumps with the ability to form a gradient of a mixture of different agents. In the case of reversed-phase LC, the stationary phase is nonpolar, separating analytes by hydrophobicity over the course of a gradient of increasing nonpolar mobile phase. The LC system is coupled to the mass spectrometer by electrospray (ES) ionization via an emitter (6). Glass or steel needles are commonly connected to the column. Particle packed capillaries for chromatography can also be used for ES without being coupled to an additional emitter (7–9). These basic attributes are shared by most LC-MS systems and differences are mainly defined by operational flow. Nano-flow LC operates at flow rates of several hundred nanoliters per minute and is the standard in proteomics due to the high sensitivity obtainable.

High flow rates in the µl to ml range, applied to columns with large inner diameters, are typically used in high-throughput or industrial-scale analysis as well as analytical MS application areas. Although these micro-flow systems limit sensitivity, recent work has demonstrated robust and reproducible performance (10,11). Reproducibility and stability of those systems are high, but drawbacks are lowered sensitivity and a need for high sample amounts. Compared to developments in sample preparation, MS instrumentation, scan modes and software, the LC apparatus has been largely unchanged in cutting edge MS-based proteomics. While identifications in proteomics experiments have doubled in single-shot experiments this can mainly be traced to improvement on the MS instrumentation and software (12–17). Current trends in LC developments aim rather towards systems for higher throughput and increasing robustness required for clinical applications (18), whereas the race for better separation in single-shot high performance runs with increasingly higher pump pressures has been comparatively abandoned. Consequently, a typically used setup for maximum sensitivity and performance for most experiments still consists of columns around 75 µm ID with a length of 20 to 50 cm, packed with sub 2 µm particles. Although, better performance could be reached by longer columns or smaller particles, both conditions would result in higher operational pressures which tend to make the LC systems unstable (4,19). For example, very high pressures can lead to leaks in the LC flow paths, resulting in poor reproducibility and subsequently a loss of measurement time.

Commercially available capillary columns in the aforementioned dimensions are expensive, especially considering how frequently they must be replaced. Therefore, many high-throughput laboratories produce packed capillaries in-house. Empty glass capillaries, ready to be packed and used, can either be purchased or produced from cheap polyimide coated capillaries using a laser puller. Typically, a gas pressure system is deployed to pack such columns with particles in the low µm range and instructions on the manufacturing process can be found online with open access (https://proteomicsresource.washington.edu/docs/protocols05/Packing_Capillary_Columns.pdf). However, this process is inherently slow and interesting methods have recently been established with the aim of speeding up the packing process with high-pressure (20) or dense bead-slurry, as in the FlashPack method (21).

Combining these principles, we here present a high-pressure packing system for capillary columns using a high-concentration bead slurry that has previously been described as beneficial for column performance (22). These high slurry concentrations and packing pressures of 1000 -3000 bar allow us to achieve packing times for 50 cm columns in the minute range with our system, compared to hours for traditional procedures. Deploying a manifold system and a pump capable of high flow rates further multiplexes packing to up to 10 columns, making column production 40 to 800 times more time efficient compared to previous systems. We observe consistently good column performance for packing pressures at over 1000 bar with no adverse effects on the column backpressure and lifetime, while packing times continued to decrease at higher pressures. We provide a detailed blueprint of the system so it can readily be set up in interested laboratories (Suppl. Table 1).

## Experimental Procedures

### Preparation of fused silica

Fused silica from Polymicro® (TSP075365 for 75 µm ID, TSP100365 for 100 µm ID or TSP150365 for 150 µm ID) was cut to 140 cm. Polyimide coating was removed by a Bunsen burner and polishing with an ethanol-soaked tissue in the middle of the cut capillary at a width of 2 cm. An electro spray emitter tip was pulled with a laser puller (Sutter P2000) at the polished part of the capillary resulting in two empty capillary columns ready to be packed.

### Sample preparation: Protein digestion and in-StageTip purification

HeLa cells were cultured in high glucose DMEM with 10% fetal bovine serum and 1% penicillin/streptomycin (all from Life Technologies, Inc.). Cells were counted using a countess cell counter (Invitrogen), and aliquots of 1×10^6^ cells were washed twice with PBS (Life Technologies, Inc.), snap-frozen and stored at −80°C. Sample preparation was carried out with the PreOmics iST kit (www.preomics.de). We used one HeLa pellet with 1 million cells per cartridge, determinate the peptide concentration after peptide cleanup via NanoDrop and adjusted the peptide concentration to 0.2 mg/ml.

### Ultra-high-pressure liquid chromatography and mass spectrometry

Samples were measured using LC-MS instrumentation consisting of an EASY-nLC 1200 ultra-high-pressure system (Thermo Fisher Scientific), coupled to an Orbitrap Exploris 480 instrument (Thermo Fisher Scientific) using a nano-electrospray ion source (Thermo Fisher Scientific). Purified peptides were separated on high-pressure packed columns as described in the results section. For each LC-MS/MS analysis with 75 µm ID columns, 500 ng peptides were used. For 100 µm ID columns, 888 ng peptides were used and for 150 µm ID columns 2000 ng peptides were used to adjust for the higher column volume.

Peptides were loaded in buffer A* (2% acetonitrile (v/v), 0.1% trifluoroacetic acid (v/v)) and eluted with a linear 105 min gradient of 5-30% of buffer B (0.1% formic acid, 80% (v/v) acetonitrile), followed by a 10 min increase to 95% of buffer B, followed by a 5 min wash of 95% buffer B. For the 75 µm ID columns flow rate was 300 nl/min, 535 nl/min for 100 µm ID columns and 1200 nl/min for 150 µm ID columns to adjust for the linear flow velocity. Column temperature was kept at 60 °C by an in-house-developed oven containing a Peltier element, and parameters were monitored in real time by the SprayQC software. MS data was acquired with a Top15 data-dependent MS/MS scan method. MS1 AGC Target was set to 300% in the 300-1650 m/z range with a maximum injection time of 25 ms and a resolution of 60,000 at m/z 200. Fragmentation of precursor ions was performed by higher-energy C-trap dissociation (HCD) with a normalized collision energy of 30 eV. MS/MS scans were performed at a resolution of 15,000 at m/z 200 with an AGC Target of 100% and a maximum injection time of 28 ms. Dynamic exclusion was set to 30 s to avoid repeated sequencing of identical peptides. Each column was equilibrated with two 120 min HeLa runs before the representative run for column cross-comparison.

### Data analysis

MS raw files were analyzed by MaxQuant software, version 1.6.11.0, and peptide lists were searched against the human Uniprot FASTA database. A contaminant database generated by the Andromeda search engine was configured with cysteine carbamidomethylation as a fixed modification and N-terminal acetylation and methionine oxidation as variable modifications. We set the false discovery rate (FDR) to 0.01 for protein and peptide levels with a minimum length of 7 amino acids for peptides and the FDR was determined by searching a reverse database. Enzyme specificity was set as C-terminal to arginine and lysine as expected using trypsin and LysC as proteases. A maximum of two missed cleavages were allowed. Peptide identification was performed with an initial precursor mass deviation up to 7 ppm and a fragment mass deviation of 20 ppm. All proteins and peptides matching to the reversed database were filtered out.

### Bioinformatics analysis

Bioinformatics analyses were performed in Python (version 3.6.4.) using Numpy (1.19.2), Pandas (1.1.4), Matplotlib (3.3.2), Seaborn (0.11.0) and Scipy (1.5.2) packages.

### Experimental design and statistical rationale

The overall experimental design was focused on making different capillary columns for proteomics experiments as comparable as possible. To achieve this, statistical analysis was done from triplicate experiments for the packing time and pressure performance experiments. Experimental conditions for column cross-comparisons were chosen to eliminate outer influences, including measurements on the similar LC and MS system and equilibration procedures.

## Results and discussion

### A high-pressure packing chamber for high density bead-slurries

A central challenge of nano-flow chromatography in proteomics laboratories is the constant demand for new capillary columns. Due to their costs, commercial columns cannot be treated as a disposable item. However, in our hands, we frequently observe highest performance only for a short lifespan for ultra-high-performance applications. Therefore, to reach the needed quantity and cost requirements, we and many other laboratories produce their own capillary columns. However, the throughput of production is limited, especially for columns with small inner diameter and extended length such as the 50 cm 75 µm inner diameter columns used in most applications in our laboratories. We produce pulled or fritted capillaries and pack them with solid phase material, typically sub 2 µm C18 beads. A skilled person can pull hundreds of empty columns within a day and fritted columns are also easy to produce. However, the packing process is an inherently low-throughput and error-prone process, which makes high-performance columns prized items in mass spectrometry laboratories. Particularly achieving longer columns length is – in our experience – a precondition for ultra-high-performance.

We hypothesized that high-throughput packing of capillary columns could be achieved by highly concentrated bead-slurries (21) in combination with very high-pressure packing (>1000 bar) (20). However, increased packing pressure and bead-slurry concentration can lead to column blocking, slowing down and eventually halting the packing procedure. Chloroform as a bead-solvent was reported as an approach to avoid this issue, because it can solvate higher bead concentrations. However, in combination with our bead-particles, we observed poor chromatographic performance during proteomic experiments. Instead, we combined elevated packing pressure with the FlashPack system (21), which prohibited bead aggregation at the column entrance via stirring.

To test our concept, we constructed a custom–made chamber for high-pressure packing, where the pressure derives from a conventional HPLC system (EASY-LC 1000 in our case). The device consists of a central chamber, containing the bead slurry and magnetic stirring bar, and has three openings. A large-bore access allows filling the chamber with the bead-slurry, a micro-bore fitting holds the capillary entrance into the chamber and a nano-viper connection is used as an inlet for the pressure from the HPLC system (Suppl. Fig. 1). The prototype packing chamber enabled us to fill single capillaries within minutes using the HPLC high-pressure pumps (950 bar). However, this system was not suited for high-throughput column production and the low pump volume of the HPLC system resulted in a non-continuous packing as the pump had to be refilled several times until a column was filled with beads.

**Fig. 1:**
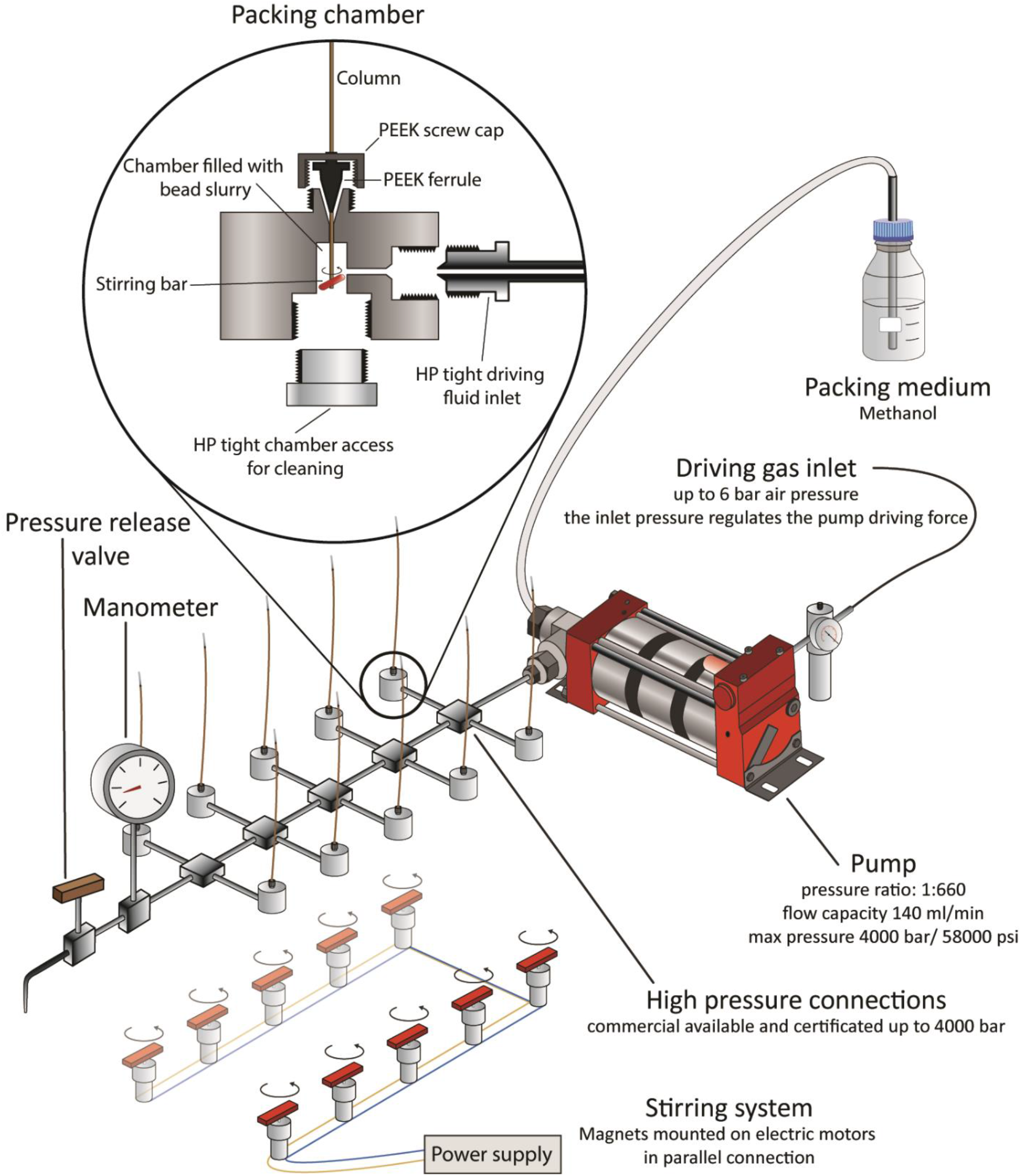
High-pressure packing station. Scheme of the high-pressure packing station with detailed description of the crucial parts. The high-pressure pump is powered by a driving gas inlet and increases the pressure of a packing medium that is provided in a large volume flask by 660-fold. The compressed packing medium is channeled to ten packing chambers, placed on top of a magnetic stirring rack. A manometer is installed to monitor the system pressure as well as a pressure release valve to facilitate time efficient system depressurization. The inset depicts a packing chamber in detail, including high-pressure fittings, stirring bar and capillary column.

Encouraged by aspects of our newly devised packing system, we set out to further streamline column production. We replaced the small volume HPLC pump with a Maximator HD-pump (Experimental Methods). This high-flow continuous system converts driving gas from a standard laboratory gas supply line at a pressure ratio of 1:660 to a fluid outlet with a maximal pressure rating of 4000 bar and maximal flow capacity of 140 ml/min (Fig 1). To use the FlashPack principle we used methanol as the packing medium, which settles C18 beads at the chamber bottom. The high flow capacity allowed us to implement multiple pump outlets for multiplex packing of up to ten columns with our station. We redesigned the original packing chamber to fit high-pressure connections (Suppl. Fig. 2). For optimal stirring, we further created a rack system with magnets mounted on electric motors via 3D printed components to fit directly underneath the packing stations (Detailed in Experimental Methods and Suppl. Fig. 3). Moreover, we connected a high-pressure range manometer to monitor packing pressure and added a pressure relief valve for efficient and controlled depressurization of the system, a notoriously time-consuming process. Even though the system is typically running at 1500 bar in our laboratory, the relief of pressure takes only 60 seconds, without flow-back from the running beads from the capillary. Additionally, the system is secured from capacity exceeding driving gas pressure by a control valve, which prevents the pump to be exposed to higher input than 6 bar. As with conventional packing systems, the weakest connection is the sealing of the capillary to the high-pressure chamber. We used a standard polyether-ether-ketone (PEEK) ferrule employed in HPLC applications in combination with a newly designed, reinforced PEEK screw cap (Suppl. Fig. 2D) to pin the column under very high pressure. Nevertheless, if the system pressure exceeds the durability of the material, the column is ejected. Due to the low compression capabilities of methanol, this is dangerous if one has body parts directly above the fitting when a rupture occurs and this must be prevented. Compared to gas, which can compress much more than liquid, no explosion risk should arise from our new packing station. Nevertheless, our recommendation is to use this device only within a secured area.

**Fig. 2:**
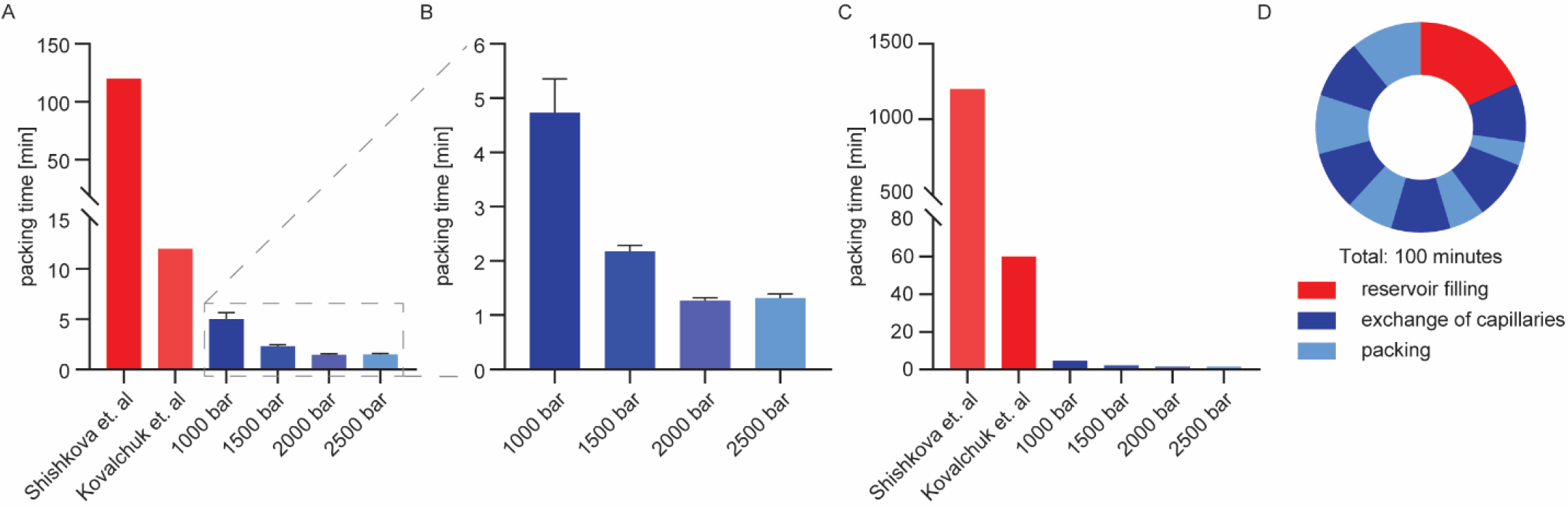
Comparison of packing times. A Packing times of single columns as described in previous efforts and for different packing pressures (data collected in triplicates, displayed with standard deviation) with a detailed view of the tested pressure conditions (B). C Production time for 10 columns considering multiplexing (2x multiplexing for Kovalchuk *et al*. and 10x for the system presented here) (20,21). D Times of a packing cycle of 10 x 5 columns, taking a total of 100 minutes with filling of the reservoir and changing of capillaries between the actual packing steps.

**Fig. 3:**
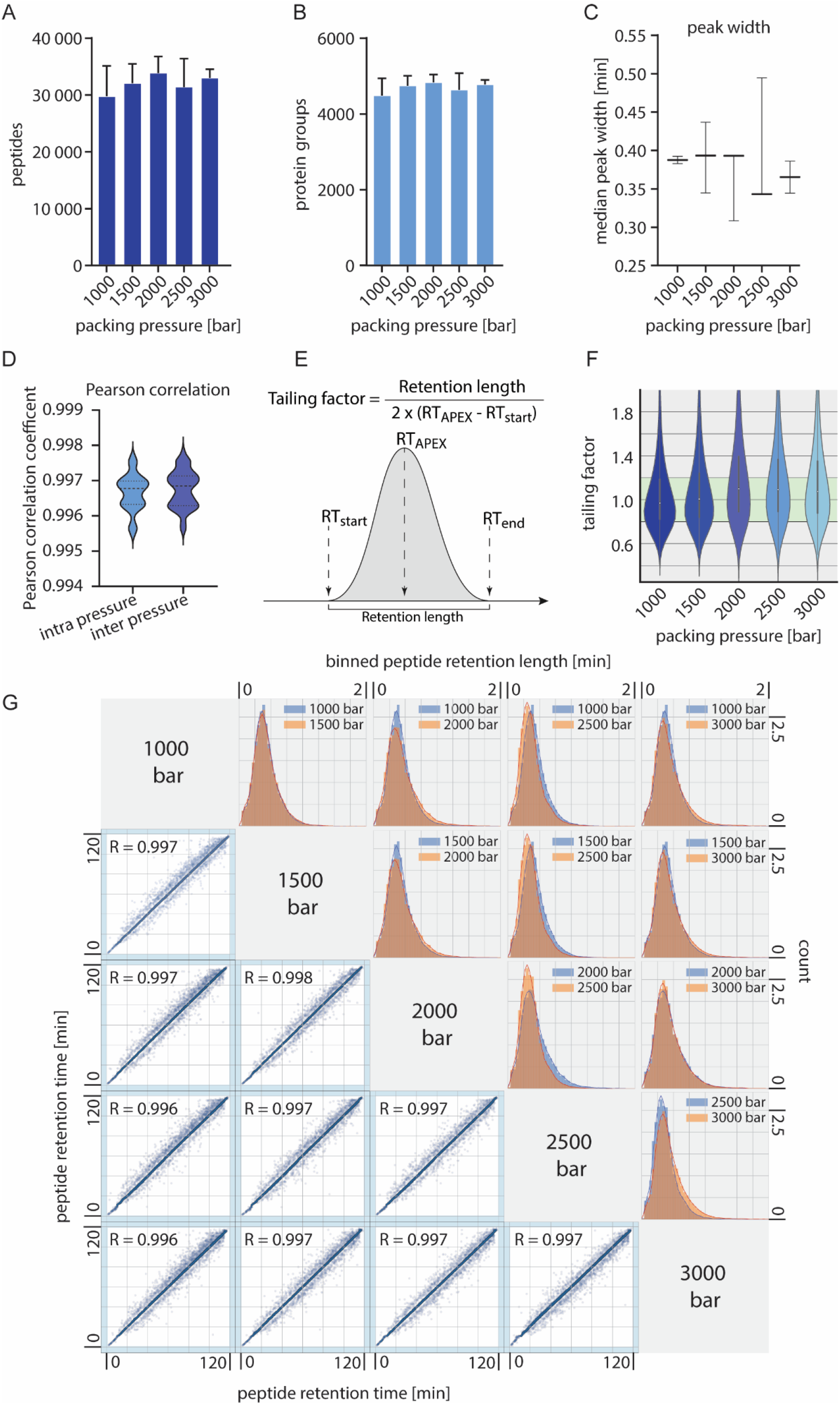
Comparison of capillary columns packed at different pressures. A, Numbers of identified peptides of triplicate measurements of 500 ng HeLa digests on columns filled at the indicated packing pressures. Peptides were separated on 50 cm, 75 µm ID columns packed with 1.9 µm Reprosil AQ Beads (Dr. Maisch) with a 2-hour gradient. B, Numbers of identified protein groups of the same conditions as in (A). Error bars indicate the standard deviation from triplicate measurements. C, Median peak widths of identified peptides. D, Distribution of Pearson correlation coefficients calculated on peptide retention times between columns packed at the same pressure and columns packed at different pressures (p-value of unpaired t-test for difference: 0.6). E, Visualization of the tailing factor calculation. F, Tailing factors for all identified peptides from runs with 75 µm ID columns and different packing pressures. G, Correlation of peptide retention times across packing conditions. The density of peptides is color-coded. The histograms show the peak width distribution of four representative runs.

### Ultra-fast column packing

The time required to fill a capillary column with beads depends on two variables, the bead concentration of the packing slurry and the flow rate through the capillary. Empty capillaries with pulled electrospray emitter have high flow rates in the µl/min range even for conventional gas-based packing bombs with lower pressure (<100 bar). However, as the bead bed grows, the flow rate through the column decreases drastically. Hence, the high-density bead slurry of FlashPack enables short packing times especially for shorter columns (21). We anticipated that combining this principle with the potentially high flow rates of our extremely high-pressure system would significantly reduce packing times.

To quantify the production throughput of our system, we consecutively packed 50 cm capillaries with 75 µm ID at different pressures (1000-2500 bar) and measured the time required. With a freshly filled bead reservoir, packing at the lowest tested pressure took on average 4.7 min. Increasing pressure to 2000 bar results in packing times just over a minute. Even higher pressure did not result in faster packing. Overall our system decreased the time for making a single column 10-to 100-fold compared to previous packing procedures (20,21) (Fig. 2A-B). Of note, the total production throughput is even higher due to multiplex-packing and the option to quickly exchange capillaries and bead slurries. This results in a 40-800 times faster column production (Fig. 2C). Once filled with bead-slurry and mounted on the high-pressure system, the packing chambers can be used to pack several columns consecutively. This merely requires depressurizing the system via the pressure relief valve and exchange the filled columns with empty capillaries. Consecutive packing of several columns from the same reservoir will decrease the packing speed due to the removal of beads from the reservoir. To fully restore packing speed, the bead chamber has to be opened and refilled, which takes about 10 min for all ten chambers together. Typically, we refilled the reservoir after five capillary exchanges. The average turn-around cycle for producing ten columns is thus 20 minutes, allowing the production of hundreds of columns in a working day (Fig. 2D). An additional advantage of the high-throughput system is that it allows us to discard non-properly packed columns, which occur in approximately 10% of cases.

The high-pressure system faces the same two main challenges as other packing stations, which are particle clogging within the capillary and bead aggregation at the column entrance. Particle clogging can only be avoided by clean working conditions. This means dust free storage and clean cutting of fused silica and the use of filtered fluids and dust free particles for bead slurry preparation. Bead aggregation from dense slurry can be circumvented by optimized stirring conditions according to the FlashPack principle (21).

### Influence of packing pressure on column performance

To evaluate the effect of packing pressure on column performance on realistic samples, we analyzed three of our laboratory standard HeLa digests on each column. Across all packing conditions, we observed no significant variation in the number of identified peptides and protein groups (Fig. 3A/B). Moreover, the median peak widths of identified peptides were comparable for all conditions (Fig. 3C). Correlation between the non-corrected retention times of peptides analyzed using columns produced at varying pressures was remarkably high (Pearson correlation coefficients > 0.996) and not significantly altered from replicates packed with similar pressure conditions (Fig. 3D).

Another factor often used to characterize column performance is the tailing factor which can be calculated as depicted in figure 3E (23). Usually, the peak width at 5% peak height is used for peak width calculation but in proteomics experiments where tens of thousands of peaks are investigated, the base-to-base peak width is typically calculated, although full width at half maximum (FWHM) is also often reported. In general, the distribution of peak shapes was wider than what would be expected from an analysis run of few analytes, but the median typically centered around the optimum of 1. The median of the tailing factor was below 1.0 for the lower and shifts above 1.0 for higher packing pressures up to a median of 1.2 (Fig 3F). In the literature tailing factors in the range between 1 – 1.2 are often described (24). The shift towards this range with the higher packing pressures could result from denser compressed bead bed. As described above the general performance was not altered for the proteomics metrics, which leads us to the conclusion that the minor change in tailing factors with higher packing pressures is not changing the LC-MS performance. This manifests in an only slightly altered distribution of peak widths between representative experiments of columns packed at different pressures (Fig. 3G). From the correlation of peptide retention times, it is visible that for all representative comparisons, the peptides elute in a narrow and reproducible time window that is not influenced by the applied packing pressure. This retention time stability is accompanied by similar separation properties of the different columns, which can be visualized directly by the retention length of analyzed molecules. Figure 3G shows bulk analysis of all identified peptides with nearly overlapping retention length distributions whereas the minor differences do not constitute a significant trend towards a better performance for lower or higher packing pressures of capillary columns. Based on these results it seems that the packing pressure has no or only minimal effect on the column performance.

### LC-MS performance of columns with different length and inner diameter

Length and inner diameter of capillary columns allow their adaptation to a plethora of sample materials and LC systems, specifically regarding separation power and backpressure. In MS-based proteomics, 75 µm ID columns in combination with flow rates in the range of 200-400 nanoliter per minute are typical. Hence, we packed such capillary columns with different lengths (20, 30, 50 cm) with our high-pressure system and compared their performance. Packing time for the shorter columns was even faster and in the range of 30 sec. The longest columns produced the smallest peak widths and subsequently resulted in the highest numbers of identified peptides and proteins (Fig. 4A-B). Interestingly, the distribution of peptide intensities did not change significantly, and the tailing factor also remained unaffected (Fig 4C-D). Over the last years the demand for high-throughput analysis has become apparent for the analysis of clinical samples, especially blood plasma as we have described before (25). This has been addressed by a novel HPLC principle with pre-formed gradients and slightly higher flow rates (18) and by higher-flow systems operating in the upper microliter per minute range (10,26). As these strategies require columns with higher inner diameter to maintain acceptable pressure during analysis, we produced columns with 75 µm, 100 µm and 150 µm ID and tested their performance.

**Fig. 4:**
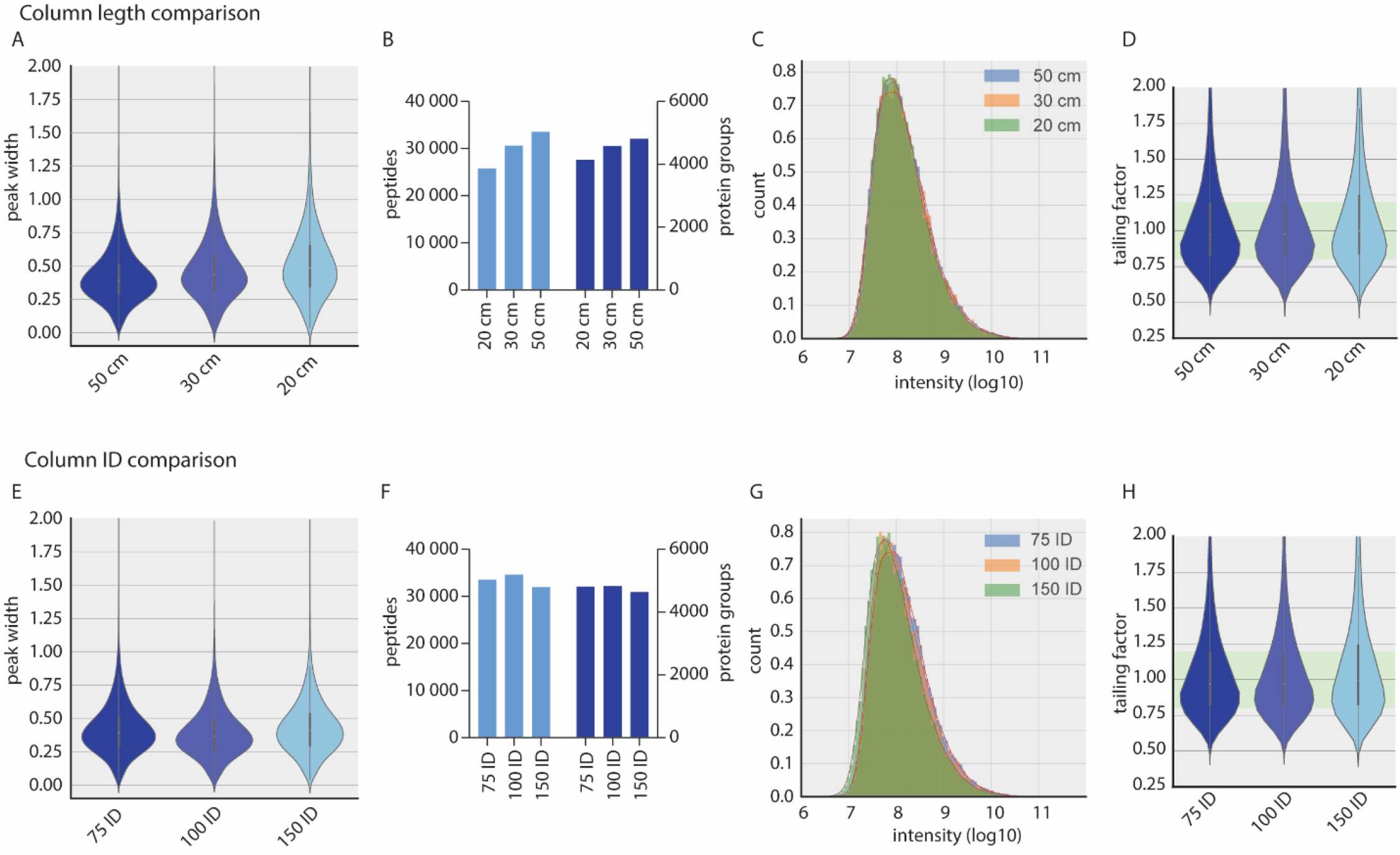
Length and inner diameter comparison. All columns were packed with 1000 bar packing pressure. A, Peak width distribution from HeLa runs with different column length with the respective number of peptide and protein identifications (B), peptide intensity distribution (log10) (C) and tailing factor distribution (D). E, Peak width distribution from HeLa runs with different column IDs with the respective number of peptide and protein identifications (F), peptide intensity distribution (log10) (G) and tailing factor distribution (H).

When comparing column inner diameters, the experimental setup has to be adapted to the conditions. To enable direct comparison of capillaries with different ID, we scaled the flow rates to reach the same linear velocities and the amount of input material to the column volume (Experimental Procedures). For the 100 µm ID columns this results in a flow rate of 535 nl/min and 888 ng of peptides for loading, whereas for the 150 µm ID column 1200 nl/min and 2 µg of peptide material was loaded to be comparable to the 300 nl/min and 500 ng employed for the 75 µm ID columns. This requirement of higher sample amount already limits the applicability of larger column diameters for samples with limited accessibility. The 1400 µl pump volume of the Easy-LC 1200 used for the experiment were sufficient to run a 2-hour gradient with the 150 µm ID column, but longer gradients or higher flow rates would exceed the capabilities of the LC-system and require higher flow rates. The larger column IDs led to slightly broader peak widths, but peptide and protein identifications were not affected. Due to the correction of the sample input amount, we have not seen a difference in peptide intensity distributions, and the peak tailing has not been affected by the column ID (Fig 4E-H).

## Conclusion

Here, we aimed to increase the throughput and to streamline the production of capillary columns for MS-based proteomics. We provide a detailed list for commercial parts and blueprints describing the construction of our high-pressure packing station. The setup can be built at relatively low costs (<$10,000), compared to the cumulative expenses for high performing commercial columns. We designed this new station to fill multiple columns simultaneously within a few minutes, which accelerates the packing process of capillary columns more than 100-fold compared to traditional gas pressure driven stations. In this way, we hope our system helps researchers streamlining the often work-intensive and fragile column production process. In addition, the extreme high pressures enable the packing of long, high-performing columns (> 50 cm). The ability to produce high-performing columns at high-throughput opens up the possibility of using capillary columns always at the peak of their performance, replacing them as soon as peak-broadening or decreased ionization is observed. Reassuring in terms of the robustness of the packing process itself and the stability achieved at exceedingly high pressures, we have not observed variation in the performance characteristics over a wide range of packing pressure from 1000 to 3000 bar. We hope the technology described here will enable laboratories of any scale to mass-produce high performance long capillary columns.

## Acknowledgments

We thank all members of the Proteomics and Signal Transduction Group at the Max Planck Institute of Biochemistry and the Clinical Proteomics Group of the NNF Center for Protein Research for help and discussions and in particular Igor Paron, Christian Deiml for technical assistance, Mario Oroshi for help with the online resource. We thank the mechanical workshop and the educational workshop especially Martin Wied, Andreas Kucher and Harry Spangenberg of the Max Planck Institute of Biochemistry for the fabrication and iterative optimization of all self-constructed parts.

## Author contributions

J.B.M-R. designed and assembled the packing station parts and carried out the bioinformatics analyses. J.B.M-R., L.S., F.H., P.G., M.M. and P.V.T. designed the experiments, performed and interpreted the MS-based proteomic analyses, generated text and figures and wrote the manuscript. M.M. supervised and guided the project, designed the experiments, interpreted MS-based proteomics data.

## Supplementary Figures

**Suppl. Figure 1.**
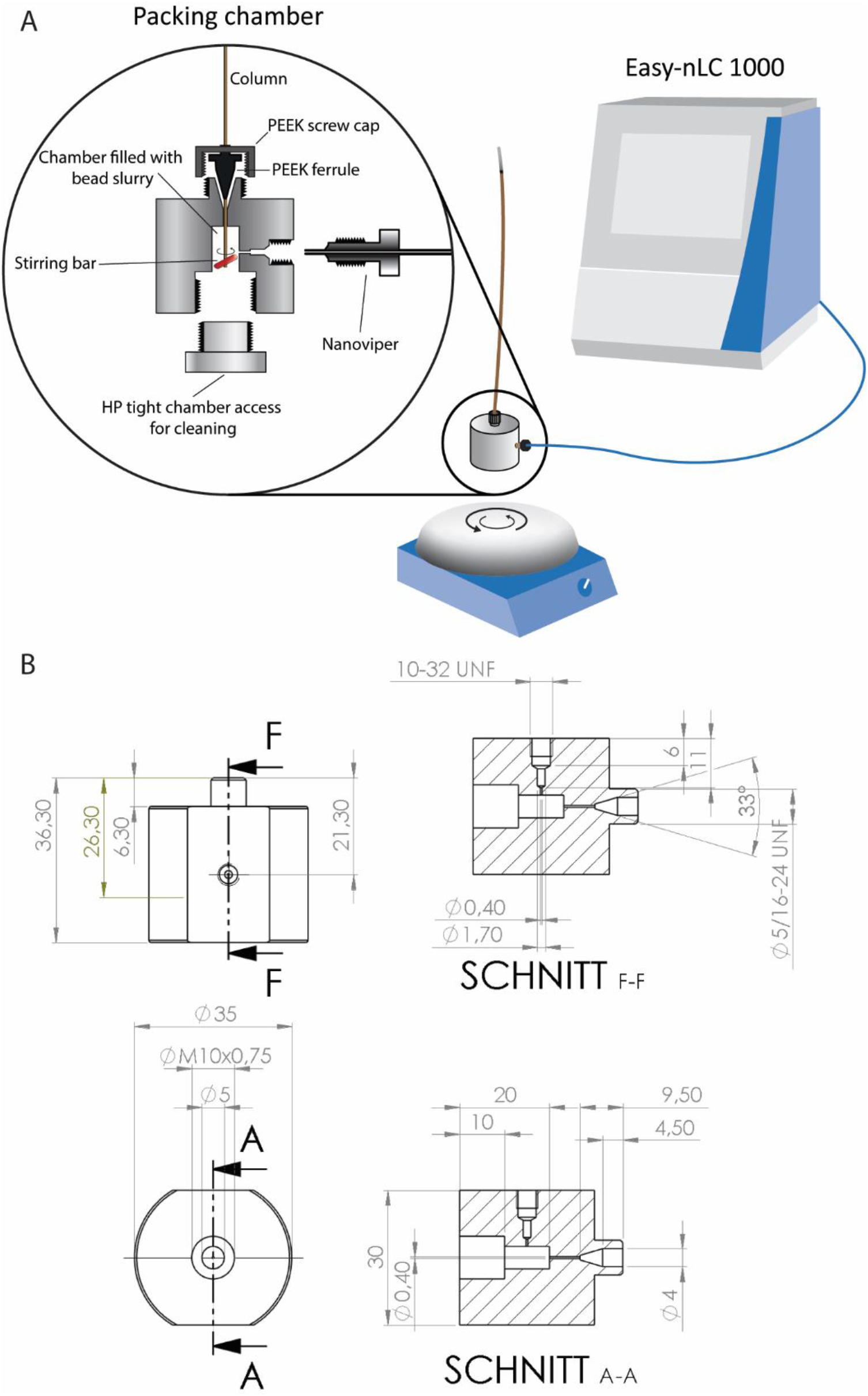
Prototype one of the high-pressure packing chamber. A, An early prototype of the packing chamber for use with a LC system with nano-viper connection output. The station was used on top of a magnetic stirrer and supplied with driving fluid and pressure from an Easy-nLC 1000. **B**, Technical drawing of the standalone prototype.

**Suppl. Figure 2.**
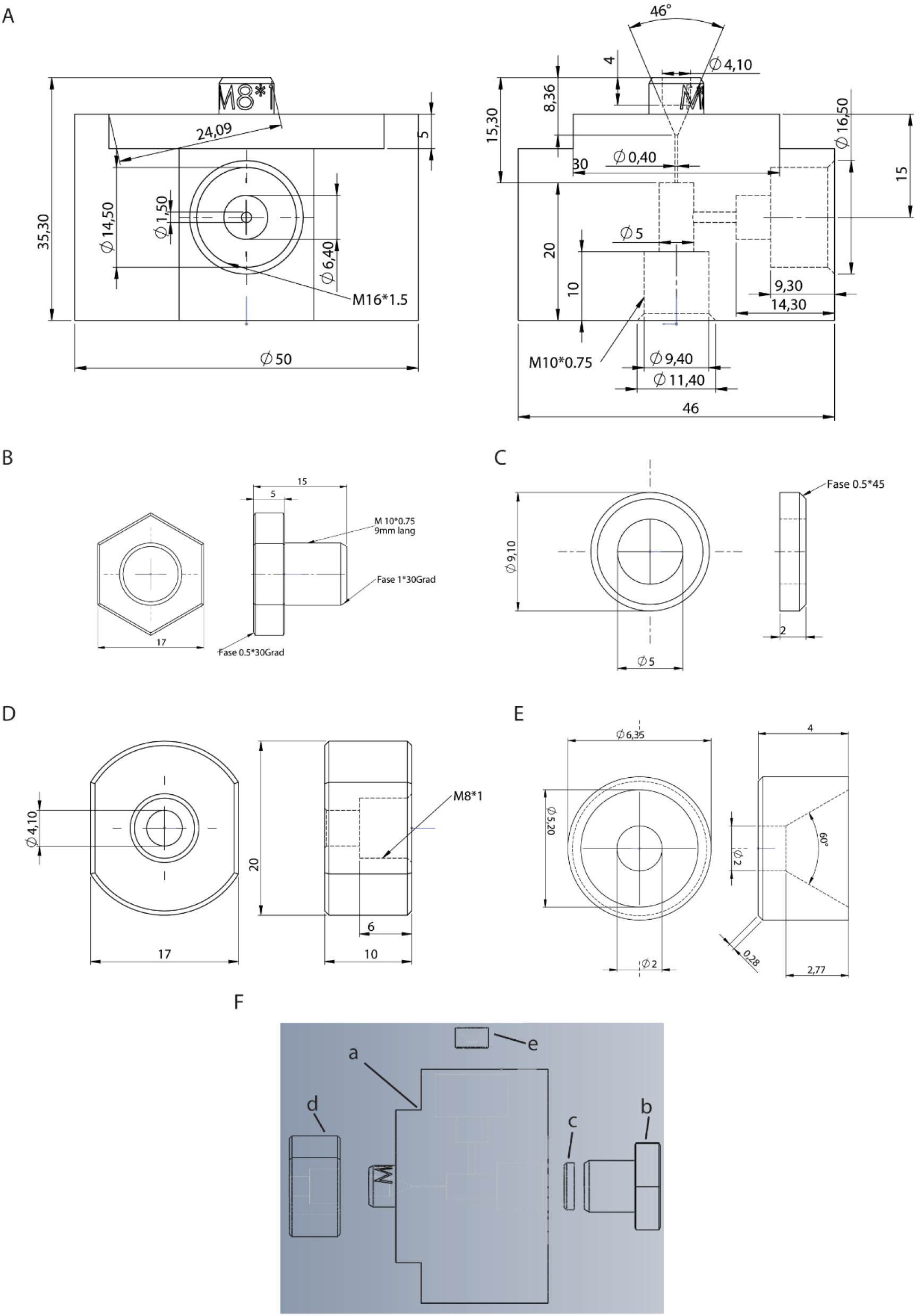
Technical drawings of the packing chamber. A, Packing chamber made from stainless steel. B, Closing screw for packing chamber and prototype one (Suppl. Fig.1) made from stainless steel. C, PEEK seal ring for the closing screw. D, GF-PEEK connection screw cap to press the PEEK ferrule into the coned fitting. E, PEEK seal ring for the connection to the HP supply system. F, Assembly of the parts A-E.

**Suppl. Figure 3.**
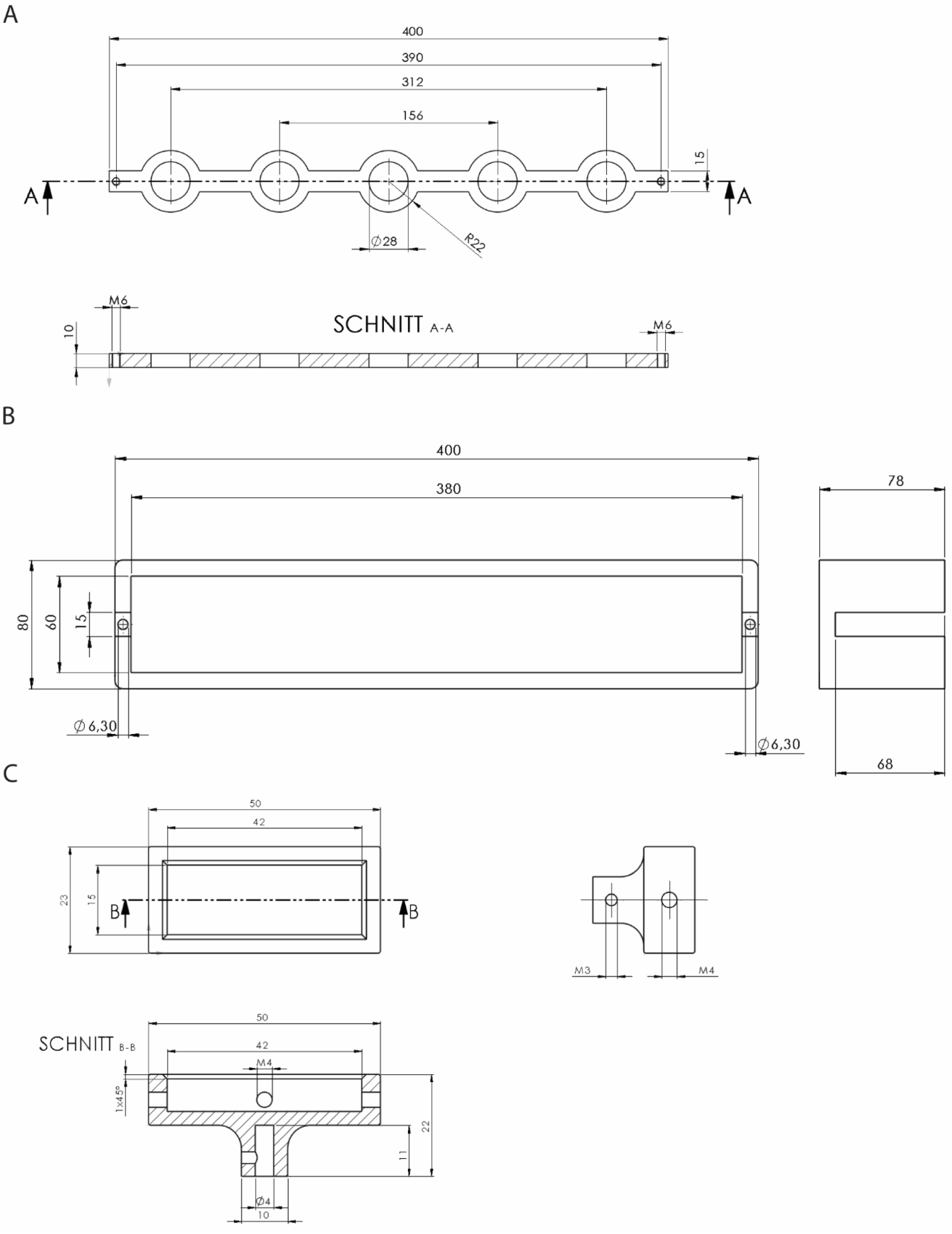
Technical drawings of the stirring system. A) Frame for the electric motors. **B**) Hight adjustable stand for the magnetic holder frame in A. C) Magnet holder to be placed on the magnetic motors 3D-printed from carbon.

